# Viruses rule over adaptation in conserved human proteins

**DOI:** 10.1101/555060

**Authors:** David Castellano, Lawrence H. Uricchio, Kasper Munch, David Enard

## Abstract

Adaptive evolution often involves fast-evolving proteins, and the fastest-evolving proteins in primates include antiviral proteins engaged in an arms race with viruses ^1-3^. Even though fast-evolving antiviral proteins are the most studied cases of primate host adaptation against viruses, viruses predominantly interact with host proteins that are broadly conserved between distant species in order to complete their replication cycle ^4^. Broadly conserved proteins are generally viewed as playing a negligible role in adaptive evolution. Here, we used a dataset of ~4,500 human proteins known to physically interact with viruses (VIPs for Virus-Interacting Proteins), to test the involvement of broadly conserved proteins in adaptive evolution against viruses. We found that VIPs conserved between animals and fungi have experienced not only high rates of adaption, but also strong adaptive events. Broadly conserved proteins that do not interact with viruses experienced very little adaptation. As a result, the arms race with viruses explains more than 75% of adaptation in the most phylogenetically conserved subset of the human proteome. Our results imply that broadly conserved proteins have played a significant role in adaptation, and that viruses were likely one of very few selective pressures that were able to force the conserved, central pillars of host cellular functions to adapt during evolution.

---

The arms race between viruses and their hosts has repeatedly involved young, fast-evolving antiviral proteins with some of the highest rates of adaptive evolution observed in the proteome. In primates, well-studied examples notably include members of the APOBEC family such as APOBEC3G ^1,3^, or members of the TRIM family such as TRIM5a ^2^. Both proteins have experienced both weak purifying selection and recurrent adaptation during primate evolution and are thought to have evolved very rapidly as a result. APOBEC3G is primate-specific, while TRIM5α is found in primates and a minority of other mammals ^5^. While most studies have focused on young, fast evolving proteins typically thought of as the main targets for adaptation, individual examples exist of more broadly conserved VIPs with substantial evidence for adaptation. For example, the antiviral factor PKR/EIF2AK2 has evolved adaptively in a large number of mammals ^6,7^ (Fig. 1A), but we found that it is nevertheless conserved well beyond mammals with clear orthologs in multiple fish genomes according to Ensembl ^5^. Moreover, we found that adaptation in PKR has occurred despite strong purifying selection, as shown by low nonsynonymous to synonymous polymorphism ratios (Methods) in both humans ^8^ and other great apes (chimpanzee,gorilla and orangutan, see Methods) ^4,9^. The example of PKR thus shows that abundant adaptation can occur even under the substantial evolutionary constraint that is characteristic of broadly conserved proteins.

**Figure 1.**
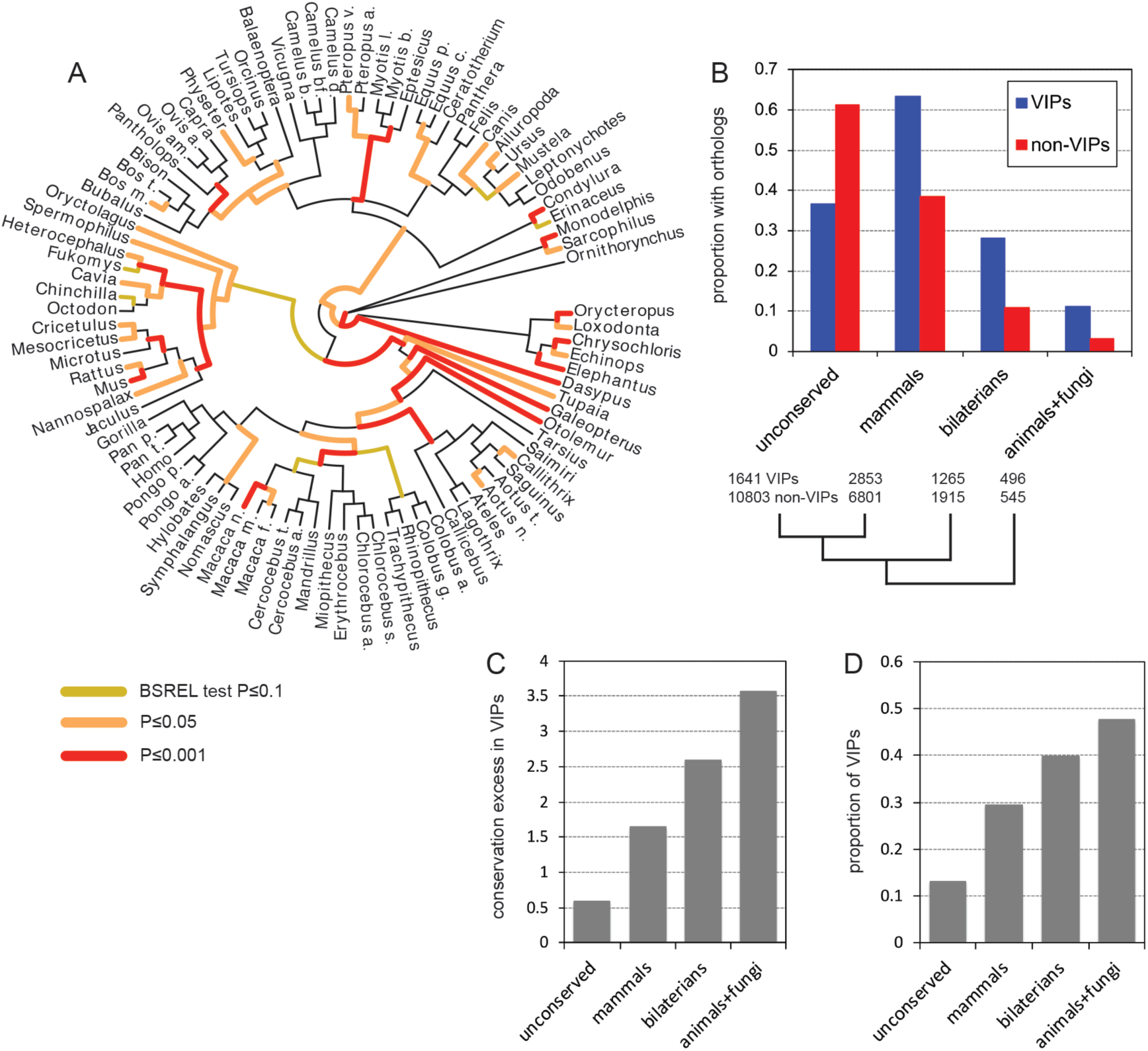
VIPs are broadly conserved across phylogenetic clades. A. Tree of 93 mammals with branches highlighted for adaptive evolution in the PKR/EIF2AK2 coding sequence. The different colors indicates the significativity of the results of the BSREL test for positive selection ^6^. Yellow indicates a weakly significant signal of positive selection, while red indicates a strongly significant signal. B. Proportions of VIPs (blue) and non-VIPs (red) that are not conserved across mammals (noted unconserved), or conserved across mammals, bilaterians or between animals and fungi (Methods). All proportions are strongly significantly different between VIPs and non-VIPs (proportion comparison test P<10^−16^). C. Relative excess of conservation at different phylogenetic breadths of VIPs compared to non-VIPs (significantly different from 1 in all cases, P<10^−16^). D. Proportions of genes conserved at different phylogenetic breadths that are VIPs. All proportions are significantly different from each other (P<10^−16^ in all cases).

Fast-evolving antiviral factors with a limited phylogenetic range are overall not representative of the thousands of host VIPs ^4^. We recently assembled a dataset of ~4,500 VIPs ^10^ that together interact with more than 20 different human-infecting viruses (supplementary table 1). VIPs experience stronger purifying selection than non-VIPs (proteins not known to interact with viruses), as shown by a lower average non-synonymous over synonymous polymorphism ratio, both in African populations ^8^ (1.14 vs. 1.39, simple permutation test P<10^−9^) and in great apes ^9^ (0.62 vs. 0.84, P<10^−9^, Methods). Furthermore, VIPs are conserved across broader phylogenetic distances than non-VIPs (Fig. 1B,C,D), with 63% of VIPs having clear orthologs conserved across mammals ^4^, versus only 39% for non-VIPs (proportion comparison test P<10^−16^). Human VIPs are also 3.6 times more likely to have a yeast Saccharomyces cerevisiae ortholog in Ensembl Compara than non-VIPs ^5^ (Fig. 1B,C, 11% vs. 3%, P<10^−16^). An elevated proportion of human-yeast orthologs are VIPs (Fig. 1D), 47% versus only 20% of human protein-coding genes overall. Conversely, VIPs are depleted in younger genes not broadly conserved across mammals compared to non-VIPs (Fig. 1B,C,D 37% vs. 61%, P<10^−16^).

To investigate if adaptation is stifled in broadly conserved VIPs and only involved younger, phylogenetically restricted VIPs, we used modern variants of the classic McDonald-Kreitman test ^11^ (MK test) to quantify adaptation in human evolution since divergence with chimpanzee (Methods). Unlike other approaches also aimed at detecting adaptation over millions of years of evolution, recent versions of the MK test control well for purifying selection and do not suffer from a decrease of statistical power in more conserved proteins ^4^. We used two recent versions of the MK test, polyDFE ^12^ and ABC-MK ^13^ (Methods), to first ask if VIPs had an overall higher rate of adaptation due to viruses compared to non-VIPs - we apply both approaches because they have complementary strengths, so evidence of adaptation obtained by both methods is stronger than either method in isolation. In the main text we put forward the results of polyDFE but also provided the ABC-MK results in Figures 2 and 3 (Methods). We simultaneously controlled for many potential confounding factors using a bootstrap test with multiple target averages (Methods). Note that we excluded immune genes from all subsequent analyses due to concerns related to the potential confounding effect of balancing selection on the MK test ^4,14^ (Methods). The conclusions of our analysis are therefore limited to non-immune genes, and in particular conserved non-immune genes (see below). In agreement with viruses driving elevated rates of adaptation in VIPs, 37% of amino-acid substitutions in VIPs were adaptive during human evolution versus 25% in non-VIPs (bootstrap test P=0.004 for polyDFE, P<10^−3^ for ABC-MK; the percentage of amino acid substitutions that were adaptive is noted a). We further found that VIPs not only experienced more adaptation, but also stronger adaptation (selection coefficient s higher than 1%; see Methods) with at least three times more strong adaptation than in non-VIPs (α=14% vs. 5%, P<10^−3^). We found similar results with ABC-MK, although the overall estimated rate of adaptation was lower (22% in VIPs vs. 8% in control non-VIPs, P<10^−3^, 16% vs. 2% for strong adaptation, P<10^− 3^). Note that lower overall estimates are expected with ABC-MK compared to polyDFE (Methods).

**Figure 2.**
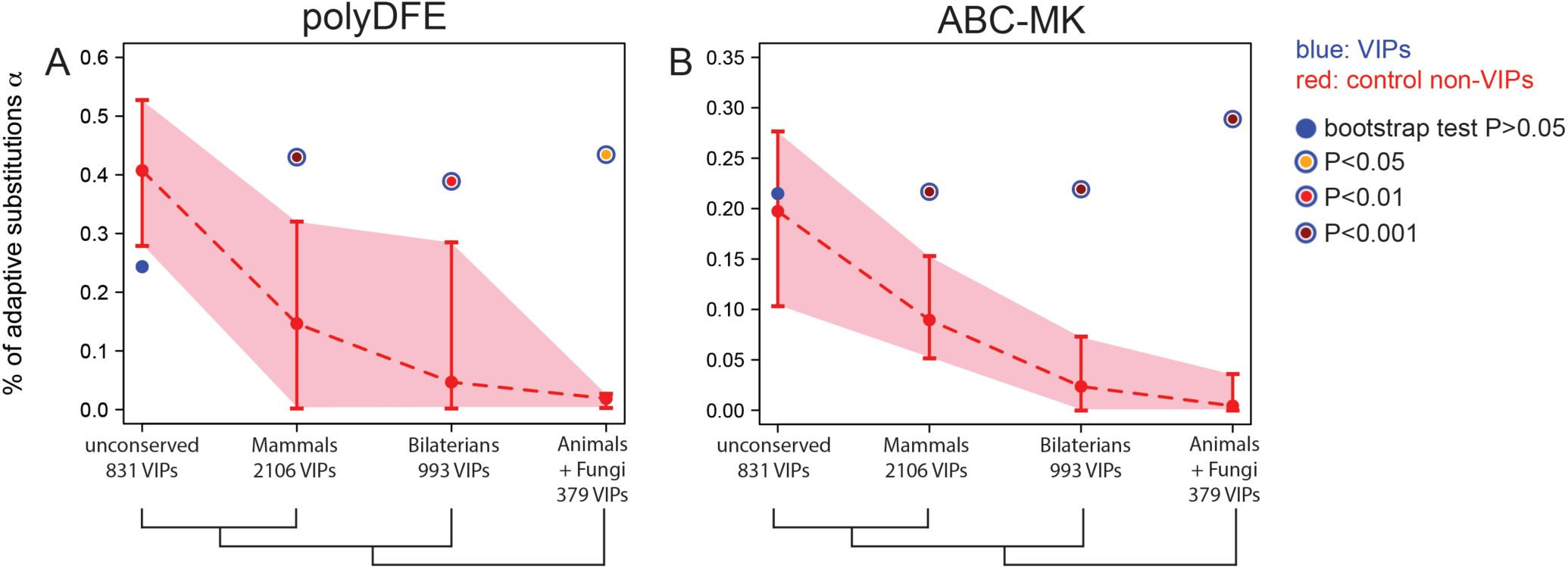
Adaptation in VIPs and control non-VIPs with increasing breadth of phylogenetic conservation. The red areas highlight the 95% confidence intervals for control non-VIPs, estimated from 1,000 iterations during the bootstrap test (Methods). Blue dots indicate α for VIPs. The legend in the figure provides the scale for how significant the bootstrap test is. A. Results for polyDFE. B. Results for ABC-MK.

**Figure 3.**
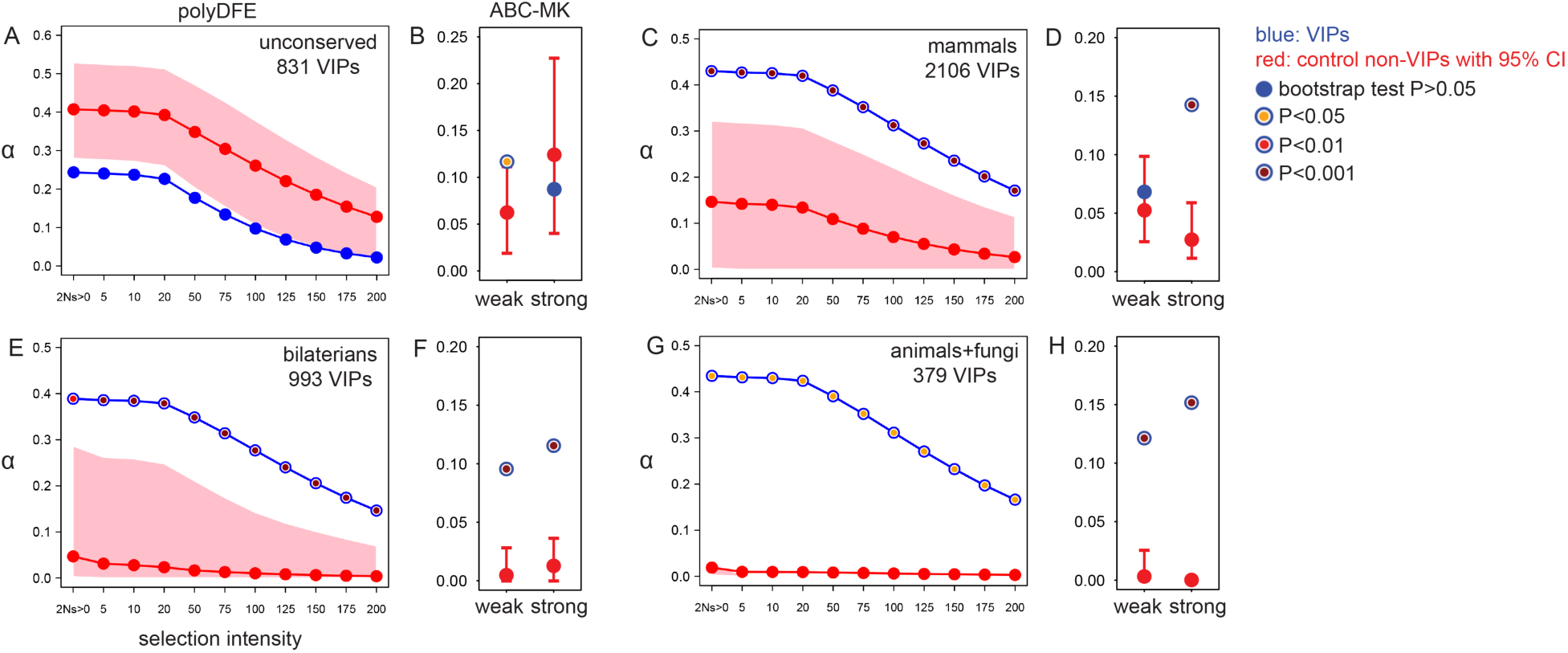
Weak and strong adaptation in VIPs compared to control non-VIPs at different breadth of phylogenetic conservation. The figure represents the estimated a for polyDFE (A,C,E and G) as a function of the intensity of selection 2Ns where N is the effective population size and s is the selection coefficient of advantageous mutations (Methods). Unlike polydFE, ABC-MK (B,D,F and H) does not provide a range of intensity but instead provides two estimates of α, one for weakly advantageous mutations of the order of 2Ns=10, and one for strongly advantageous mutations of the order of 2Ns=100 or greater ^13^ (Methods). Red areas represent the 95% confidence intervals for matched control non-VIPs. A and B. Genes not conserved across mammals, noted unconserved. C and D. Genes conserved across mammals. E and F. Genes conserved across bilaterians. G and H. Genes conserved between animals and fungi.

We then asked how the increase in adaptation was distributed across VIPs with different levels of phylogenetic orthologous conservation ^5,15^ (Figure 2). Unexpectedly, we found no increase and even a decrease of adaptation at VIPs compared to non-VIPs when restricting the comparison to younger genes that do not have broadly conserved orthologs across mammals (Methods; Fig. 2A,B and 3A,B). This may be due to the fact that the rate of adaptation is elevated in these genes overall, including in non-VIPs (Fig. 2A,B and 3A,B). By contrast, we found much higher adaptation in VIPs compared to non-VIPs with broader phylogenetic conservation. We estimated α =43% in VIPs that are conserved in mammals, versus 15% in similarly conserved control non-VIPs (Fig. 2A,B and 3C,D). Adaptation in VIPs was also much stronger, with α=17% for strong adaptation (s>1%) in VIPs versus 3% for non-VIPs (Fig. 3C,D). The estimated α remained at 39% in VIPs conserved across bilaterians, and at 15% for strong adaptation (Fig. 2A,B and 3E,F). Adaptation was very low in similarly conserved control non-VIPs, with α=5% overall and 0.4% for strong adaptation (Fig. 2A,B and 3E,F). Remarkably, α still remained at 43% overall and 17% for strong adaptation in VIPs shared between animals and fungi while being estimated even closer to zero at 2% overall (0.3% for strong adaptation) in similarly conserved control non-VIPs (Fig. 2A,B and 3G,H). These results were robust to the choice of method used to annotate orthologous conservation ^5,15^ (Extended Data Fig. 1), and suggest that the rate of adaptation at VIPs remains stable across levels of phylogenetic conservation, while at the same time higher levels of conservation have strongly limited adaptation in non-VIPs. Furthermore, broadly conserved proteins that interact with multiple viruses experienced strongly increased adaptation, as expected if viruses were indeed the driving force behind these patterns (Extended Data Fig. 2). Again, the results obtained with ABC-MK were broadly concordant with polyDFE (Figs. 2 and 3).

Broadly conserved proteins therefore experienced sharply different patterns of evolution, depending on whether they interact with viruses or not. VIPs have experienced substantial adaptation, and particularly strong adaptive events. An important consequence is that the majority of human adaptation in broadly conserved proteins, and especially strong adaptation, was likely driven by viruses. We quantified the proportion of all adaptive amino acid changes that were driven by viruses at different breadths of conservation (Methods). In proteins conserved in mammals, we estimated that viruses drove 28% of all adaptation (Methods), 55% for proteins conserved in bilaterians, and 95% for proteins conserved between animals and fungi (25%, 56% and 78% for ABC-MK estimates, respectively). For strong adaptation (s>1%), we estimated that 40% of strong adaptive events may have been driven by viruses in proteins conserved across mammals, 85% in proteins conserved across bilaterians and 98% for proteins conserved across animals and fungi (45%, 64% and 84% for ABC-MK, respectively). Other types of pathogens such as bacteria or eukaryotic pathogens have not supported a similar level of impact on conserved proteins ^16^ (SI). Therefore, viruses have likely ruled over adaptation in the conserved human proteome. Interestingly, different viruses (supplementary table 1) had similar contributions to adaptation in broadly conserved proteins, and no single virus that we tested represented and unusually large proportion of adaptive events (SI).

Together, these results paint an unexpected picture of proteome-wide adaptation against viruses that cannot be inferred from generalizing the known examples of young, fast-evolving antiviral proteins ^1-3^. Our analyses show that viruses do not seem to increase overall adaptation in non-immune, younger proteins - in contrast, viruses drive the vast majority of strong adaptation events in broadly conserved proteins. However, because they are specialized in restricting viruses, fast-evolving, immune antiviral proteins can accumulate many protective adaptive substitutions, and thus undoubtedly represent highly significant players in the arms race with viruses. This nonetheless raises the question of why viruses, if they represent such a strong selective pressure, do not increase overall adaptation in younger non-immune VIPs. One possible explanation is that young VIPs have not interacted with very many viruses during evolution. However, younger VIPs interact with significantly, but only slightly fewer viruses on average compared to VIPs conserved in mammals (1.73 vs. 1.91 respectively, simple permutation test P<10^−3^). Another possible explanation is that viruses are only one of many other selective pressures experienced by younger proteins, as shown by the high levels of adaptation in younger non-VIPs. We also cannot fully exclude that an increase in adaptation in younger VIPs might be masked by a hypothetical dampening effect of frequent balancing selection on the MK test (Methods). This, however, would not change our main finding that the majority of human adaptation in broadly conserved, functionally central proteins is strong adaptation driven primarily by viruses. More generally, our results challenge the view that conserved proteins do not adapt, highlight the importance of strong adaptive events in otherwise conserved proteins, and provide a more complete and more quantitative picture of genome-wide adaptation against viruses.

## Methods

### VIPs dataset

The dataset of VIPs has been extensively described ^4,10^. Supplementary table 1 provides the complete list of VIPs. The table also provides which VIPs have orthologs across mammals, bilaterians or animals and fungi. We used a total of 4,494 VIPs that had stable Ensembl IDs (supplementary table 1) between Ensembl v83 and Ensembl v73. Of these 4,494 VIPs, we could use 3,838 that included all the confounding factors we controlled for (supplementary table 2; see the rest of Methods). We excluded genes (including 1,092 immune VIPs, supplementary table 1) annotated as immune by the Gene Ontology (http://www.qeneontoloqv.org/. Annotations GO:0006952, GO:0006955 or GO:0002376) due to concerns regarding potential effects of balancing selection on the MK test (See Methods, Balancing selection and the MK test).

### Constraint in PKR/EIF2AK2

We measured level of constraint in PKR/EIF2AK2 as the ratio of non-synonymous polymorphism over synonymous polymorphism both in both humans ^8^ and other great apes (chimpanzee,gorilla and orangutan) ^4,9^. In humans we used polymorphism in African populations from the 1000 Genomes Project phase 3 ^8^. In great apes, we summed the non-synonymous polymorphism over chimpanzee, gorilla and orangutan populations included in the Great Apes Genome Project ^4,9^, and did the same for synonymous polymorphism. More specifically we used the ratio of the number of non-synonymous variants over the number of synonymous variants plus one to avoid null denominators (noted pN/(pS+1)).

In human African populations, for PKR/EIF2AK2 pN/(pS+1)=5/12=0.42 vs. 1.27 for the proteome average. This represents the 46^th^ lowest ratio out of 496 coding sequences with the same number of variants (pN+pS). In great apes, pN/(pS+1)=3/14=0.21 vs. 0.91 for the proteome average. This is the 71^st^ lowest ratio out of 476 coding sequences with the same number of variants. In summary, the low pN/(pS+1) ratio in both humans and other great apes shows that PKR/EIF2AK2 has experienced both strong constraint and intense adaptation.

### Divergence and polymorphism data

The divergence and polymorphism data used for this study have already been described in detail in Uricchio et al. ^13^. In brief, we used human-chimpanzee-orangutan alignments of 17,740 orthologous coding sequences to identify fixed substitutions on the human branch. For polymorphism we used the variants present in the 661 individuals of the 1000 Genomes Project African populations phase 3 ^8^, and their derived allele frequencies measured across Africa. Supplementary table 3 provides all the data necessary to run the MK test.

### Using polyDFE and ABC-MK to account for the strength of selection

In order to verify that our results were not sensitive to a specific version of the MK test, we used two implementations of the MK test that use different approaches, namely polyDFE ^12^ and the ABC-MK test ^13^. The polyDFE implementation of the MK test uses a maximum likelihood approach, while ABC-MK uses Approximate Bayesian Computation to estimate α, the proportion of adaptive amino acid changes that were adaptive in a given evolutionary lineage. Compared to other existing methods, both polyDFE and ABC-MK do not make the simplifying assumption that advantageous mutations do not contribute to polymorphism. This assumption can be problematic when a substantial proportion of adaptation is weak adaptation, with adaptive mutations taking a longer time to go to fixation, as may be the case in human evolution ^13^. Both polyDFE and ABC-MK account for the contribution of weakly advantageous mutations and use this contribution to estimate separate rates of weakly and strongly advantageous substitutions. This ability to distinguish between weak and strong adaptation is important to estimate the overall α accurately. It is also important for this specific study, to determine whether a specific α in conserved coding sequences reflects only weak adaptation with limited fitness implications, or instead strong adaptation with important fitness implications and potentially substantial functional consequences. Segregating weakly advantageous mutations are explicitly integrated in the model used by polyDFE given that polyDFE estimates the full distribution of fitness effects, while in ABC-MK it is the shape of the α-curve as a function of derived allele frequency that is determined by the abundance and frequency of segregating weakly advantageous variants.

In addition to distinguishing between weakly and strongly advantageous substitutions, polyDFE and ABC-MK each have other complementary advantages. Indeed, polyDFE has the advantage that it can estimate α solely from polymorphism data without requiring divergence information. This is important because estimates of α can lose accuracy due to long-term changes in population sizes ^17^. For example, current estimates of modern human effective population sizes are smaller than the older effective population sizes that have been estimated for earlier human evolution, closer to the split with chimpanzee ^18-20^. This means that modern humans have likely experienced weaker purifying selection compared to earlier human evolution. The issue then is that the current number of non-synonymous variants is higher than the long-term number of non-synonymous variants, due to more effectively neutral deleterious variants segregating and even sometimes fixing. Such variants would have been eliminated under earlier higher population sizes. At the same time, the long-term number of non-synonymous variants available for fixation, not the higher current number, determines the number of fixed non-synonymous substitutions. If not taken into account, the currently higher number of non-synonymous variants then results in underestimating α in human evolution ^13^. By avoiding using divergence altogether, polyDFE also avoids this potential discrepancy between the current and the long-term numbers of non-synonymous variants. Note however that the fact that polyDFE uses only polymorphism implies that the estimated **a** are only representative of more recent human evolution.

ABC-MK has the advantage of accounting for the effect of linked background selection on rates of adaptation, and especially on rates of weak adaptation. Indeed, polyDFE uses measures of the strength of background selection ^21,22^ at different locations in the genome to estimate how much it reduced the fixation of weakly advantageous variants. We can thus use ABC-MK to directly account for the potential confounding effect of background selection. Note that we also account for background selection when using polyDFE, by matching sets of VIPs with control sets of non-VIPs with similar levels of background selection ^22^ in the bootstrap test. For this latter reason we put forward the results of polyDFE in the main text and provided the results of ABC-MK in Figures 2 and 3 (see also below). Note that ABC-MK is sensitive to long term fluctuations in population size.

The ABC-MK test was run as in Uricchio et al ^13^. In order to achieve reasonable computing times to convergence (typically less than 48 hours per run for the top 1% longest runs), we ran polyDFE with the flexible, default model C and with subsets of 20 randomly sampled individuals out of the 661 African individuals in the 1000 Genomes Project phase 3. We also used the basin hopping option (500 maximum iterations) of polyDFE to improve convergence, as well as the grouping option with the following groups: [1,2] [3,20] [21,30] [31,40]. These groups were defined to capture the differences in the shape of the Site Frequency Spectrum induced by the presence or absence of segregating weakly advantageous variants ^13^.

### Understanding differences between polyDFE and ABC-MK estimates of α

By comparing the results of polyDFE with the results of ABC-MK, we were able to show that the two implementations of the MK test predict similar contributions of viruses to human protein adaptation (see main text and Methods, Estimating the contribution of viruses to total adaptation). However, polyDFE and ABC-MK estimated different α (Figs. 2 and 3). Because we used the version of polyDFE based only on polymorphism to avoid the confounding effect of ancient fluctuations in population size, the results of polyDFE reflect relatively recent human evolution. How much evolutionary time is captured by polyDFE notably depends on how long it takes for weakly advantageous mutations to increase in frequency. For very weakly advantageous mutations that would fix on average twice as fast as neutral mutations, the evolutionary time captured by polyDFE might then represent half or 4Ne generations (Ne being the effective population size), or 20,000 generations if considering Ne~10,000 in modern humans. Considering a generation time between 25 and 30 years ^23^, polyDFE would then capture the last 500,000 to 600,000 years of human adaptive evolution. Note however that this reasoning neglects the fact that even older adaptive events that brought advantageous variants to higher frequencies could still participate in the differences between the SFS of non-synonymous and synonymous variants, even long after the driving selective pressures have disappeared ^24^. Although it is thus unclear how much evolutionary time is captured by polyDFE, it is very likely to be much less than the divergence time between humans and chimpanzee.

ABC-MK uses divergence and thus reflects human evolution since divergence with chimpanzee. We therefore do not expect to estimate the same α with polyDFE and ABC-MK. The α estimates from ABC-MK may also represent underestimates (see above) due to long-term population size fluctuations, which may further explain the differences observed between polyDFE and ABC-MK. In support of this, we noticed that while polyDFE based on polymorphism estimated α close to zero (2%) in non-VIPs broadly conserved between humans and yeast, polyDFE based on polymorphism and divergence estimated a substantially negative α (-15%) for the same proteins. This suggests that the downward bias due to ancient population size fluctuations becomes apparent in the form of negative α estimates in proteins that experience very little adaptation, but where α should still be positive and very close to zero. It is interesting to note that the 17% difference (2% to -15%) between the two estimates of polyDFE used with or without divergence matches well the differences observed at all VIPs between polyDFE and ABC-MK (37% vs. 22%).

For these reasons we chose to put forward the results of polyDFE based on polymorphism only in the main text, although it represents adaptation in relatively recent human evolution. The polyDFE version of the MK test also likely better accounts for very weakly advantageous mutations compared to ABC-MK. Indeed, the ABC-MK version of the MK test assumes weakly advantageous mutations with 2Ns=10, but not lower ^13^, whereas polyDFE does not have such a limitation. This predicts that ABC-MK may underestimate α by not including very weakly advantageous mutations with 2Ns below 10. Note that it is a matter of debate whether such weakly advantageous mutations that behave like nearly neutral mutations should be considered as advantageous in the first place ^25^.

### Estimating the contribution of viruses to total adaptation

In addition to estimating a, we used a in VIPs and non-VIPs to estimate how much adaptation was driven by viruses. We defined the contribution of viruses to total protein adaptation as the percentage of all adaptive amino acid changes that were driven by viruses. In VIPs, not all adaptive amino acid changes were necessarily driven by viruses. As an estimate of the baseline adaptation in VIPs in the absence of viruses, we used the average α from control non-VIPs. The average α in control non-VIPs is indeed a good estimate of the baseline because control non-VIPs were selected to have similar genomic properties as VIPs (see next sections), interactions with viruses set aside. We then calculated the difference between α in VIPs and the baseline α provided by control non-VIPs to estimate the proportion of adaptive amino acid changes actually due to viruses in VIPs. Then, by multiplying this proportion by the total number of amino acid changes in VIPs, we were able to estimate the number of adaptive amino acid changes due to viruses in VIPs. To estimate the number of adaptive amino acid changes in non-VIPs, we estimated α for all non-VIPs at least 50kb away from VIPs to avoid clustered, undiscovered VIPs as well as linkage effects with nearby VIPs. Note that these VIPs include all VIPs far enough from VIPs, and not only those non-VIPs that were used as controls in the bootstrap test and that represent only a subset of all non-VIPs. The total number of adaptive amino acid changes in non-VIPs was then calculated as the total number of amino acid changes in all non-VIPs including those close to VIPs, multiplied by α in the non-VIPs at least 50kb away from VIPs. Finally, we could calculate the proportion of adaptation due to viruses as the number of adaptive amino acid changes due to viruses in VIPs, divided by the sum of the total number of adaptive amino acid changes in VIPs and of the same number in non-VIPs.

### Sets of conserved, orthologous proteins

For our analysis, we used only orthologs that are more likely to be conserved not only at the sequence level but also at the functional level compared to other types of homologs ^15^. Viruses tend to target conserved cellular functions for their replication ^4,26^. To ensure that we worked with human genes that not only had partial sequence conservation, but also more likely functional conservation, we only included, one to one, best reciprocal orthologs. Such orthologs are less likely to evade the selective pressure of viruses by redistributing the targeted host functions through duplication and the formation of paralogs. Adaptation in such conserved functions is therefore more likely to have to happen at the orthologous sequence level.

The different sets of conserved proteins we used are based on orthology information from a previous study of VIPs in mammals ^4^ and from the Ensembl Compara database ^5^. In a previous study we identified 9,589 human coding sequences (CDS) that are best-reciprocal homology hits between human CDS and the CDS of 23 other eutherian mammals, still have the same Ensembl Identifiers as in our previous study in Ensembl v83, and are included in the 17,740 human-chimp-orangutan orthologs that we used (supplementary table 3). These 9,589 CDS represent the set of CDS broadly conserved across mammals (Figs. 2 and 3). Of these 9,589 CDS, we used 7,681 non-immune that had information for all the confounding factors taken into account by the bootstrap test (supplementary table 1). These 7,681 CDS only include CDS with non-zero (>0.0005 cM/Mb over a 200kb window centered half-way between the genomic start and the genomic end of each gene) recombination rates to avoid gaps in the recombination map we used ^27^. These 7,681 CDS include 2,106 VIPs (supplementary table 1). The set of CDS not broadly conserved across mammals that we defined as a set of younger CDS is made of the 7,668 CDS that remain after removing the 9,589 CDS conserved across mammals from the human-chimp-orangutan orthologous CDS that we used to run the MK test on the human branch. Of these 7,668 CDS, we used 5702 non-immune with factors available for the bootstrap test (supplementary table 1). Of these 5702 CDS, 831 are VIPs (supplementary table 1).

The 3,016 CDS conserved across Bilaterians (Figs. 2 and 3) are CDS annotated as one to one orthologs between human and Drosophila melanogaster by Ensembl Compara v93. Of these 3,016 CDS, we used 2,548 non-immune that had information for all the confounding factors taken into account by the bootstrap test (supplementary table 1). Again, these 2,548 CDS only include CDS with non-zero recombination rates. Of these 2,548 CDS, 994 are VIPs (supplementary table 1). The 978 CDS conserved across animals and fungi (Figs. 2 and 3) are CDS annotated as one to one orthologs between human and Saccharomyces cerevisiae by Ensembl Compara v93. We used 808 of these 978 CDS (supplementary table 1), including 379 VIPs (supplementary table 1).

In addition to the Ensembl Compara orthologs, we verified that our results were robust to the particular annotation tools of orthologs by using predictions of orthologs from Wormhole ^15^. that provides machine learning predictions of orthologs based on the integration of 17 different ortholog prediction algorithms. Wormhole provides scores of confidence for orthology predictions ranging from zero to one, from low to high confidence. We validated the robustness of the results found with Ensembl Compara by using Wormhole best reciprocal orthologs with confidence score above 0.5 or 0.9. The number of orthologous VIPs between human and Drosophila melanogaster or between human and yeast varied substantially as a function of the confidence score (Extended Data Fig. 1). However, the estimates of adaptation in VIPs and control non-VIPs remained very similar (Extended Data Fig. 1), showing that these results are robust to the particular set of orthologs used.

### Confounding factors

An important hurdle when trying to detect the impact of a specific selective pressure by quantifying adaptation in associated genes (VIPs to detect the impact of viruses in our case) is that these genes may also vary in different other ways that also affect rates of adaptation compared to the rest of the genome. For example, VIPs are more highly expressed across many tissues than non-VIPs ^4,10,26^, and stronger expression could hypothetically lead to higher rates of adaptation in VIPs in an indirect way that has nothing to do with a causal impact of viruses. In order to account for such potential confounding factors, we adopted an exhaustive approach and included as many possible confounding factors as we could think of based on previous literature and available data. In total, we included 13 different factors that could have an impact on the estimation of α (supplementary table 2). These factors include:

-**DS**: the number of fixed synonymous substitutions in humans since divergence with chimpanzee. Controlling for DS accounts for any bias that would affect α through DS rather than through the effect of adaptation on DN, the number of non-synonymous fixed substitutions.

-**PN** and **PS**: we matched sets of VIPs with control sets of non-VIPs with the same average number of non-synonymous variants PN in African populations, and the same number of synonymous variants PS. We matched PS for the same reason as DS. We matched PN to build control sets of non-VIPs with the same average amount of strong purifying selection as VIPs. Indeed, the stronger the purifying selection, the higher the number of amino acid changing positions where mutations are not tolerated even at low frequencies.

-**GC content**. GC content correlates with many other potential confounding factors and is a good proxy for long-term recombination rate. Matching GC content thus controls for potential confounders such as biased gene conversion and long-term recombination rate ^28^. We used the GC content measured in 50kb windows centered on genes, half-way between gene starts end gene ends.

-**recombination rate**. Local recombination rate can have a profound impact on adaptation by affecting the amount of linkage with deleterious or other advantageous mutations and by affecting the levels of background selection. We therefore matched VIPs with control non-VIPs with a similar recombination rate from the genetic map of Hinch et al. ^27^. The recombination rates were estimated in 200kb windows centered half-way between the genomic start and genomic end of Ensembl protein coding genes genes. We excluded any VIP and non-VIP with a recombination rate less than 0.0005 cM/Mb from our analysis to avoid confusing genes where the recombination rate is actually close to zero, and genes where there is simply a gap in the genetic map (missing data).

-**coding sequence length**. To compare VIPs with non-VIPs that contained the same amount of information we matched coding sequence length.

-**average expression across 53 GTEx tissues (**https://gtexportal.org/home/): VIPs tend to be more strongly expressed across a broad range of tissues compared to non-VIPs ^4,10,26^. We matched VIPs with control sets of non-VIPs with the same average total cumulated expression level (in RPKM) across 53 tissues represented in GTEx V6 ^29^. We used the log2 of RPKM to account for the wide dispersion of expression values across several orders of magnitude.

-**average GTEx expression in testis**: VIPs are more expressed in whole testes, and male reproductive tissues have been shown to evolve under higher rates of adaptation ^30,31^. To avoid confusing adaptation in response to viruses with reproductive adaptation we controlled for GTEx testis expression. We used the log2 of RPKM.

-**average GTEx expression in lymphocytes**: VIPs are more expressed in GTEx lymphocytes. To avoid confusing immune adaptation in general and adaptation dues specifically to viruses, we controlled for GTEx lymphocyte expression (in addition to removing all genes annotated as immune by GO).

-**the number of protein-protein interactions**: VIPs tend to have more interacting partners than non-VIPs in the human protein-protein interaction (PPI) network, and the PPI number has been shown to affect rates of adaptation ^32^. We matched VIPs and controls for the log2 of the number of interacting partners ^32^.

-**McVicker’s B for background selection**: We matched VIPs and control non-VIPs for McVicker’s B value to account for the effect of BGS on rates of adaptation and especially weak adaptation ^22^.

-**the coding sequence density**: to further account for potential linkage effects with functional and deleterious variation in a gene’s neighborhood we matched VIPs and non-VIPs for the overall coding sequence density in a 50kb window centered half-way between the genomic start and end of each gene.

-**the overall density of PhastCons conserved elements**: to further account for linkage effects including with potential deleterious variation in non-coding conserved elements we matched VIPs and control non-VIPs for the density of PhastCons conserved elements ^33^ in 50kb windows.

### Accounting for confounding factors: the bootstrap test with multiple target averages

We created a bootstrap test called the bootstrap test with multiple target averages to match VIPs with control sets of non-VIPs. Previously we had created a bootstrap with only one target average for only one factor ^4^. For a given set of VIPs, we build a control set of non-VIPs that that has the same overall average values for all the 13 factors described above. The matching process represents a bootstrap because the same non-VIP can be resampled and represented multiple times in the control set of non-VIPs. We build the control sets of non-VIPs gradually until the number of non-VIPs matches the number of VIPs tested (counting non-VIPs sampled multiple times as many times as they appear in the control set). Each non-VIP is sampled randomly and added to the control set if the averages for the 13 factors in the growing control set all stay within matching limits defined around the VIP average. These matching limits are defined as follows. If a factor is on average higher in VIPs compared to non-VIPs as is for example the case for overall gene expression, we define a lower bound at 0.95 times the VIP average for the control set of non-VIPs. In this case we do not set a higher bound because the randomly sampled non-VIPs tend to have much lower expression than VIPs anyways. If a factor is on average lower in VIPs compared to non-VIPs as is for example the case for the number of non-synonymous variants PN, we define an upper bound at 1.05 times the VIP average for the control set of non-VIPs. In this case we do not set a lower bound because the randomly sampled non-VIPs tend to have much higher PN than VIPs. Furthermore, in the cases where not enough matching non-VIPs are found and the matching process takes too long, we slightly expand the lower and higher bounds of the permissible range. We expanded the permissible range to plus or minus 7% for the test with genes broadly conserved in animals and fungi (Figs. 2 and 3), and to plus or minus 10% for tests with VIPs that interact with two or more viruses (Extended Data Fig. 2). We sample non-VIPs as potential controls only if they are located 50kb or further from VIPs. We do this because neighboring genes tend to be functionally more similar, which means there could be a chance that non-VIPs close to VIPs might be more likely to be yet undiscovered VIPs. Although the same non-VIPs can be represented multiple times in the bootstrap control set, we set a limit that each individual non-VIP cannot represent more than 2% of the control set.

An important difficulty when gradually building control sets of non-VIPs while trying to simultaneously match 13 different factors is that only a small proportion of individual non-VIPs in the genome may actually match closely the combination of averages across 13 different factors. This can be problematic when starting to build a control set of non-VIPs, because the first non-VIPs that would result in averages close to the VIP set averages are VIPs that are themselves close to the averages. This is a problem because many more diverse combinations of non-VIPs can collectively match the VIP averages, while individually not matching these averages at all (by compensating each other). In order to ensure that we select control sets of non-VIPs that include as many non-VIPs as possible and not only the non-VIPs that happen to closely match the VIP set averages, we use a starting group of 50 “fake non-VIPs” that all individually match the VIP set averages perfectly. We then gradually add real non-VIPs to this fake set of non-VIPs. Then, even if the first added real non-VIP has factor values far from the VIP set averages, there is still a good chance that the averages of all factors for the 50 fake + 1 real non-VIPs will still be within the authorized range of averages. If the 50 fake and one real non-VIPs are within the authorized average range, we keep the real non-VIP and we remove one fake non-VIP supposed to perfectly match the VIP set average. We then verify that the average for the 49 remaining fake non-VIPs and the one real non-VIP also fall within the acceptable range. We then repeat the addition of a real non-VIP and removal of a fake non-VIP until there are no fake non-VIPs left, at which point we simply start adding non-VIPs to the 50 first real non-VIPs. This procedure ensures that the control set of non-VIPs includes non-VIPs with diverse combinations of factors’ values while still matching VIPs in terms of overall averages. In this manuscript, for each comparison of VIPs with non-VIPs, we created 1,000 sets of control non-VIPs. We also did not actually add non-VIPs to growing sets of non-VIPs one by one, but two by two, because it also increased the number of VIPs that could be used as controls due to the compensation effect between pairs of different non-VIPs whose average for confounding factors happens to fall closer to the target averages.

### Balancing selection and the MK test

Balancing selection such as frequency-dependent selection or heterozygous advantage can maintain functional variants at intermediate frequencies. Such balanced variants could potentially affect estimates of α by the MK test if they are common enough, especially at immune genes or genes interacting with pathogens in general ^14^. This would specifically happen in the case where balancing selection increases the number of non-synonymous variants at intermediate frequencies, which is expected to result in underestimating α. In a previous study ^4^, we found that α for immune VIPs (VIPs with GO immune annotations) is lower than α at immune non-VIPs. No such pattern was observed when comparing non-immune VIPs and non-immune non-VIPs. We speculated at the time that it could be due to balancing selection increasing the number of non-synonymous variants at intermediate frequencies in immune VIPs. Here, we tested this hypothesis with the new, larger set of 4,494 VIPs. To test if immune VIPs experienced balancing selection increasing the number of intermediate-frequency non-synonymous variants, we tested if they have an increased number of non-synonymous variants at derived allele frequencies between 0.4 and 0.6 (close to 0.5) compared to 1,000 control sets of non-VIPs built using the bootstrap test with multiple target averages (see above, factors same as the other tests conducted in the manuscript). Note that this comparison is conservative given that overall the number of non-synonymous variants in VIPs is lower than in non-VIPs due to stronger purifying selection in VIPs. Despite the conservativeness of the test, we did find a significantly higher number of non-synonymous variants at (78 vs. 58.7 expected on average, bootstrap test P=0.023), suggesting that balancing selection might indeed have increased non-synonymous variation at intermediate frequencies. Thus, because of the potential underestimating effect of balancing selection on α in immune VIPs but also in other immune genes ^14^, we decided as a precaution to not include immune genes in any of the analyses conducted in the manuscript. Importantly, excluding immune genes does not change the main result that adaptation in broadly conserved proteins is mostly driven by viruses. Removing immune genes is also justified by the fact that it is already very-well known and would not represent a new result that immune, antiviral VIPs have experienced abundant adaptive evolution.

## Acknowledgements

We thank Thomas Bataillon for clarifications on polyDFE. DC was funded by the Danish Research Council for Independent Research grant DFF-4181-00358.

## Author Contributions

Conceived and designed the experiments: DE, DC, and KM. Performed the experiments: DC, LHU, and DE. Wrote the paper: DE.

## Supplementary Information

**Extended data Figure 1.**
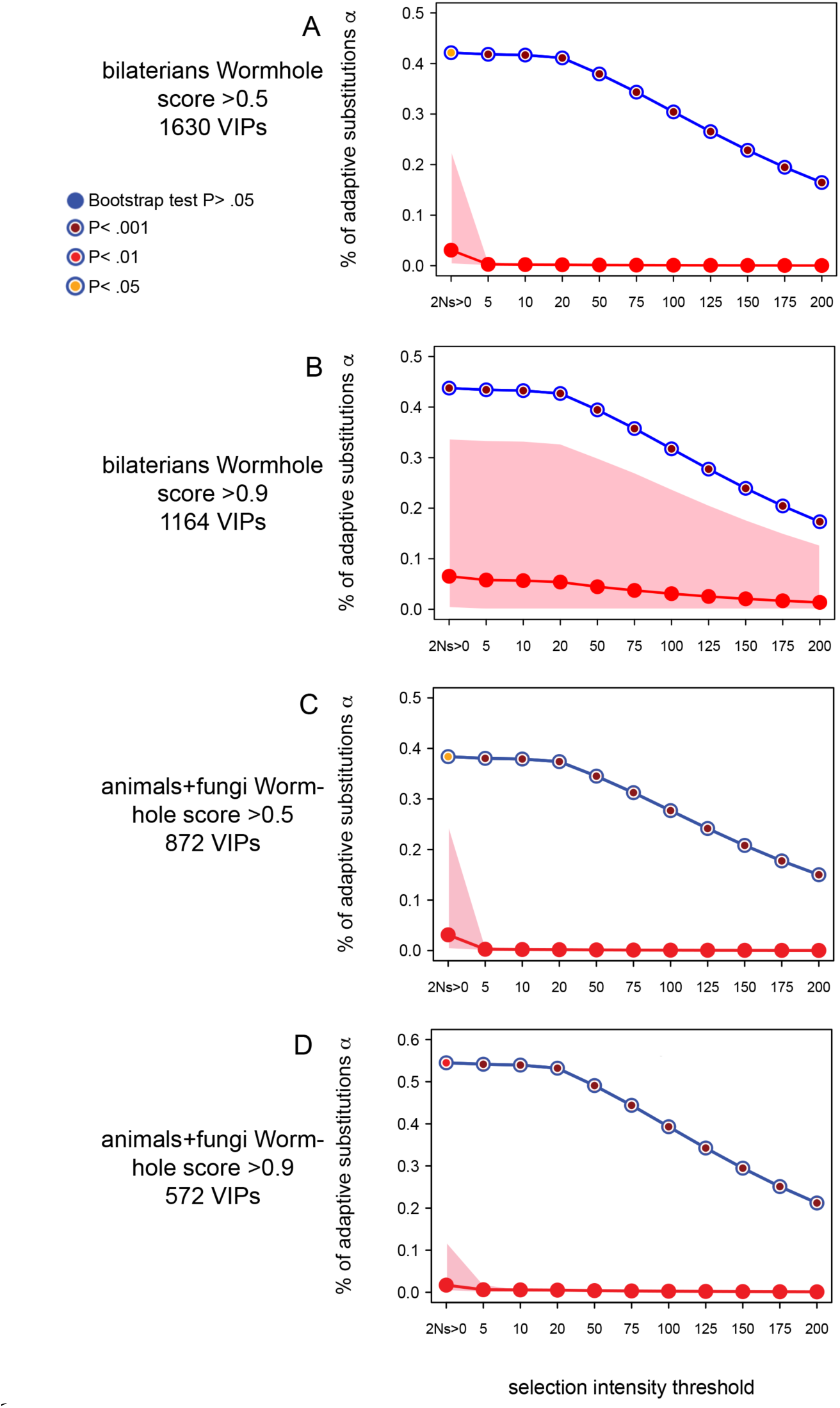
Robustness of the results to the particular annotation of conserved orthologs. Same legend as Fig. 3.

**Extended data Figure 2.**
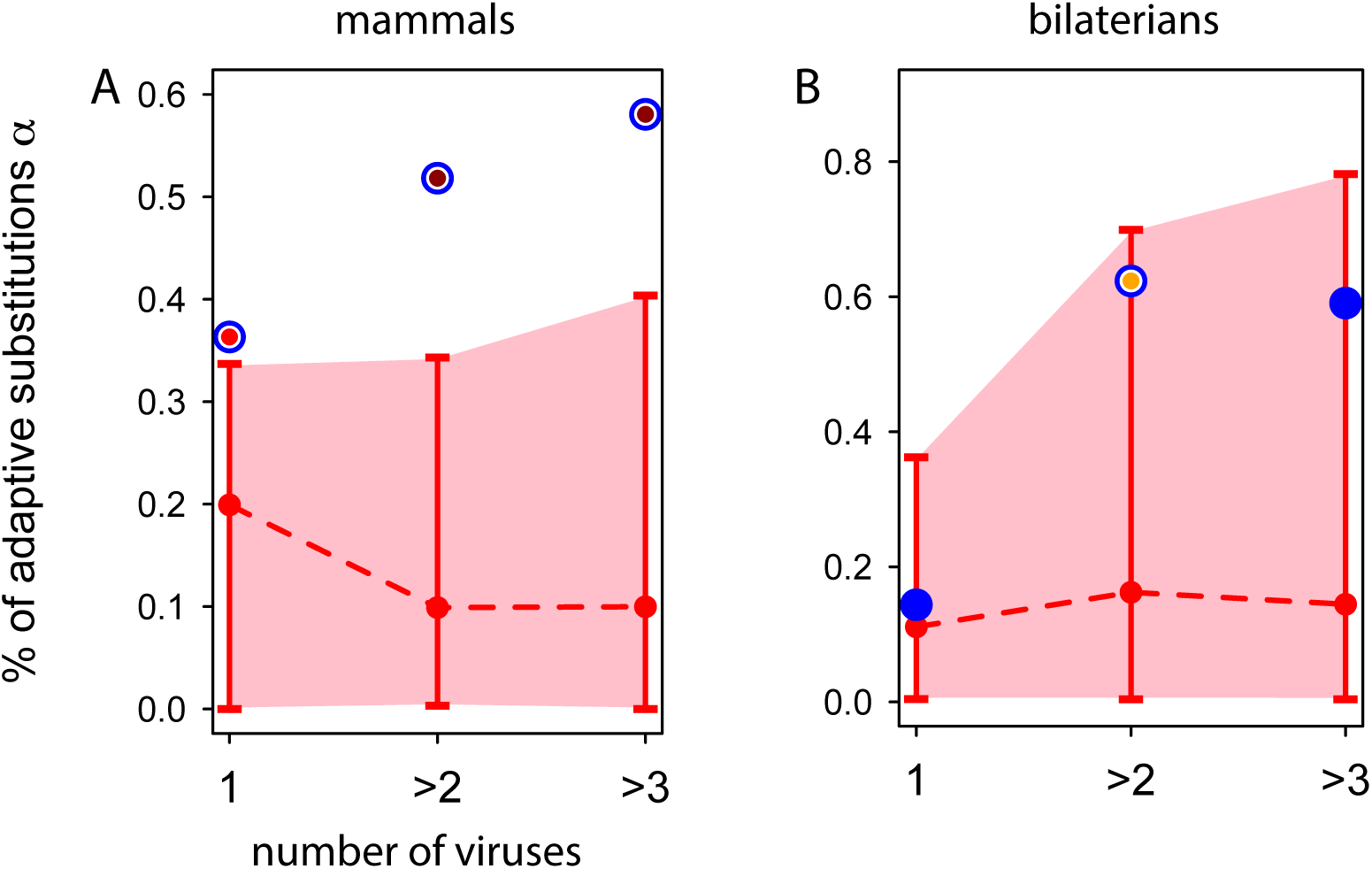
Higher rates of adaptation in VIPs that interact with multiple viruses. Legend as in Fig. 2. A. Mammals. B. Bilaterians. The number of VIPs that interact with only one virus and are broadly conserved in animals and fungi was too low to get reliable estimates, with confidence intervals for a ranging from zero to one, showing insufficient information for convergence of polyDFE.

Supplementary table 1 is provided as an excel spreadsheet.

### Testing the impact of specific viruses

We tested whether specific viruses drove more or less adaptation compared to other viruses in conserved proteins. For each of 11 viruses with more than 50 testable VIPs conserved across mammals, we used polyDFE to measure a. We then compared the estimated α for the tested virus with the estimates obtained from controls sets of VIPs that interact with other viruses. The set of VIPs for the tested virus and the control VIPs that interact with other viruses were matched for the number of viruses they interact with using the bootstrap test. This was done to avoid confusing the effect of a specific virus with the possibility that the VIPs of a specific virus might just happen to interact with more viruses than the average VIP. We further only used control VIPs that are at least 50kb away from the VIPs for the tested virus. This was done to account for the fact that neighboring genes tend to be functionally more similar and might thus be more likely to interact with the same viruses (members of a same complex, expression in the same cell types, etc.). None of the 11 viruses we tested stood out, with estimated α not significantly different from α estimated for the control VIPs of other viruses (bootstrap test P>0.05 in all cases). The same was true when using VIPs conserved across bilaterians or VIPs conserved across animals and fungi (bootstrap test P>0.05 in all cases again).

### Other types of pathogens

To test the impact of non-viral pathogens we used one dataset of physical interactions between human proteins and bacteria extracted from the Intact database and that we described and used previously ^16^. We also used a manually curated dataset of Plasmodium-interacting proteins that represents a good sample of interactions for eukaryotic pathogens infecting humans ^16^. We used the bootstrap test to ask if BIPs or PIPs (Bacteria-Interacting Proteins or Plasmodium-Interacting Proteins) were enriched for human adaptation compared to non-BIPs or non-PIPs, respectively. We ran the bootstrap test with multiple target averages the same way we did for VIPs and non-VIPs, but in addition to the 13 confounding factors that we accounted for when comparing VIPs and non-VIPs, we also controlled for the number of viruses BIPs or PIPs interact with since we already know that viruses increased adaptation in human proteins. We tested BIPs and PIPs with polyDFE at different levels of phylogenetic conservation (mammals, bilaterians, and animals and fungi). Unlike for VIPs, we found no significant enrichment of protein adaptation at BIPs, nor at PIPs (bootstrap test P>0.05 for all conservation levels). This suggests that viruses had a stronger impact on conserved protein adaptation than bacteria or eukaryotic pathogens.

## References

1. Sawyer, S. L., Emerman, M. & Malik, H. S. Ancient adaptive evolution of the primate antiviral DNA-editing enzyme APOBEC3G. PLoS Biol 2, E275, doi:10.1371/journal.pbio.0020275 (2004).

2. Sawyer, S. L., Wu, L. I., Emerman, M. & Malik, H. S. Positive selection of primate TRIM5alpha identifies a critical species-specific retroviral restriction domain. Proc Natl Acad Sci U S A 102, 2832–2837, doi:10.1073/pnas.0409853102 (2005).

3. Zhang, J. & Webb D. M. Rapid evolution of primate antiviral enzyme APOBEC3G. Hum Mol Genet 13, 1785–1791, doi:10.1093/hmg/ddh183 (2004).

4. Enard, D., Cai, L., Gwennap, C. & Petrov, D. A. Viruses are a dominant driver of protein adaptation in mammals. Elife 5, doi:10.7554/eLife.12469 (2016).

5. Zerbino, D. R. et al. Ensembl 2018. Nucleic Acids Res 46, D754–D761, doi:10.1093/nar/gkx1098 (2018).

6. Kosakovsky Pond, S. L. et al. A random effects branch-site model for detecting episodic diversifying selection. Mol Biol Evol 28, 3033–3043, doi:10.1093/molbev/msr125 (2011).

7. Elde, N. C., Child, S. J., Geballe, A. P. & Malik, H. S. Protein kinase R reveals an evolutionary model for defeating viral mimicry. Nature 457, 485–489, doi:10.1038/nature07529 (2009).

8. Genomes Project, C. et al. A global reference for human genetic variation. Nature 526, 68–74, doi:10.1038/nature15393 (2015).

9. Prado-Martinez, J. et al. Great ape genetic diversity and population history. Nature 499, 471–475, doi:10.1038/nature12228 (2013).

10. Enard, D. & Petrov, D. A. Evidence that RNA Viruses Drove Adaptive Introgression between Neanderthals and Modern Humans. Cell 175, 360–371 e313, doi:10.1016/j.cell.2018.08.034 (2018).

11. McDonald, J. H. & Kreitman, M. Adaptive protein evolution at the Adh locus in Drosophila. Nature 351, 652–654, doi:10.1038/351652a0 (1991).

12. Tataru, P., Mollion, M., Glemin, S. & Bataillon, T. Inference of Distribution of Fitness Effects and Proportion of Adaptive Substitutions from Polymorphism Data. Genetics 207, 1103–1119, doi:10.1534/genetics.117.300323 (2017).

13. Uricchio, L. H., Petrov, D. A. & Enard, D. Exploiting selection at linked sites to infer the rate and strength of adaptation. bioRxiv, doi:10.1101/427633 (2018).

14. Bitarello, B. D. et al. Signatures of Long-Term Balancing Selection in Human Genomes. Genome Biol Evol 10, 939–955, doi:10.1093/gbe/evy054 (2018).

15. Sutphin, G. L., Mahoney, J. M., Sheppard, K., Walton, D. O. & Korstanje, R. WORMHOLE: Novel Least Diverged Ortholog Prediction through Machine Learning. PLoS Comput Biol 12, e1005182, doi:10.1371/journal.pcbi.1005182 (2016).

16. Ebel, E. R., Telis, N., Venkataram, S., Petrov, D. A. & Enard D. High rate of adaptation of mammalian proteins that interact with Plasmodium and related parasites. PLoS Genet 13, e1007023, doi:10.1371/journal.pgen.1007023 (2017).

17. Rousselle, M., Mollion, M., Nabholz, B., Bataillon, T. & Galtier, N. Overestimation of the adaptive substitution rate in fluctuating populations. Biol Lett 14, doi:10.1098/rsbl.2018.0055 (2018).

18. Burgess, R. & Yang, Z. Estimation of hominoid ancestral population sizes under bayesian coalescent models incorporating mutation rate variation and sequencing errors. Mol Biol Evol 25, 1979–1994, doi:10.1093/molbev/msn148 (2008).

19. Schrago, C. G. The effective population sizes of the anthropoid ancestors of the human-chimpanzee lineage provide insights on the historical biogeography of the great apes. Mol Biol Evol 31, 37–47, doi:10.1093/molbev/mst191 (2014).

20. Takahata, N., Satta, Y. & Klein, J. Divergence time and population size in the lineage leading to modern humans. Theor Popul Biol 48, 198–221, doi:10.1006/tpbi.1995.1026 (1995).

21. Charlesworth, B., Morgan, M. T. & Charlesworth, D. The effect of deleterious mutations on neutral molecular variation. Genetics 134, 1289–1303 (1993).

22. McVicker, G., Gordon, D., Davis, C. & Green, P. Widespread genomic signatures of natural selection in hominid evolution. PLoS Genet 5, e1000471, doi:10.1371/journal.pgen.1000471 (2009).

23. Moorjani, P. et al. A genetic method for dating ancient genomes provides a direct estimate of human generation interval in the last 45,000 years. Proc Natl Acad Sci U S A 113, 5652–5657, doi:10.1073/pnas.1514696113 (2016).

24. Kimura, M. & Ohta, T. The Average Number of Generations until Fixation of a Mutant Gene in a Finite Population. Genetics 61, 763–771 (1969).

25. Galtier, N. Adaptive Protein Evolution in Animals and the Effective Population Size Hypothesis. PLoS Genet 12, e1005774, doi:10.1371/journal.pgen.1005774 (2016).

26. Halehalli, R. R. & Nagarajaram, H. A. Molecular principles of human virus protein-protein interactions. Bioinformatics 31, 1025–1033, doi:10.1093/bioinformatics/btu763 (2015).

27. Hinch, A. G. et al. The landscape of recombination in African Americans. Nature 476, 170–175, doi:10.1038/nature10336 (2011).

28. Duret, L. & Arndt, P. F. The impact of recombination on nucleotide substitutions in the human genome. PLoS Genet 4, e1000071, doi:10.1371/journal.pgen.1000071 (2008).

29. Consortium, G. T. Human genomics. The Genotype-Tissue Expression (GTEx) pilot analysis: multitissue gene regulation in humans. Science 348, 648–660, doi:10.1126/science.1262110 (2015).

30. Voight, B. F., Kudaravalli, S., Wen, X. & Pritchard, J. K. A map of recent positive selection in the human genome. PLoS Biol 4, e72, doi:10.1371/journal.pbio.0040072 (2006).

31. Nielsen, R. et al. A scan for positively selected genes in the genomes of humans and chimpanzees. PLoS Biol 3, e170, doi:10.1371/journal.pbio.0030170 (2005).

32. Luisi, P. et al. Recent positive selection has acted on genes encoding proteins with more interactions within the whole human interactome. Genome Biol Evol 7, 1141–1154, doi:10.1093/gbe/evv055 (2015).

33. Siepel, A. et al. Evolutionarily conserved elements in vertebrate, insect, worm, and yeast genomes. Genome Res 15, 1034–1050, doi:10.1101/gr.3715005 (2005).

